# Recovery from form-deprivation myopia in chicks is dependent upon the fullness and correlated colour temperature of the light spectrum

**DOI:** 10.1101/2021.04.28.441740

**Authors:** Arumugam R. Muralidharan, Low Wan Yu Shermaine, Lee Yong Chong, Veluchamy A. Barathi, Seang-Mei Saw, Dan Milea, Raymond P. Najjar

**Author notes:** Corresponding author **Correspondence and reprint requests to:** Dr. Raymond P. Najjar, Singapore Eye Research Institute, The Academia, 20 College Road, Discovery Tower Level 6, Singapore 169856.

## Abstract

**Background:** To evaluate the impact of full-spectrum light-emitting diodes (LEDs) mimicking sunlight on ocular axial elongation and refractive error development in a chicken model of myopia.

**Methods:** A total of 39 chicks (Lohmann brown), 1 day-old, were randomly distributed into 3 groups. Animals were housed for 28 days in a temperature-controlled enclosure, under a 12/12h light/dark cycle of isoluminant (∼285 Lux) fluorescent [n = 18, (4000K, FL-4000)] or Sunlike-LED [n=12, (4000K, SL-4000); n = 9, (6500K, SL-6500)] white lights. Myopia was induced monocularly in all chicks by random occlusion of one eye with a frosted diffuser, from day 1 post-hatching (D1) until D14. On D14, diffusers were removed, and recovery from myopia was monitored under the same experimental light condition. Axial length (AL), refractive status, choroidal thickness and anterior chamber depth were recorded on days 1, 7, 14, 22 and 28. *Ex vivo* scleral collagen fibre thicknesses were measured from scanning electron microscopy images. Differences in outcome measures between eyes and groups were compared using 2-way repeated-measures ANOVA.

**Results:** There was no significant difference between groups in the AL and refraction of form-deprived (FD) eyes during form-deprivation (D1 to D14). FD eyes of animals raised under SL-4000 and SL-6500 recovered more rapidly from excessive axial elongation than those of animals raised under FL-4000, by D22 and D28. Correspondingly, the refractive status of FD eyes exposed to SL-4000 and SL-6500 was close to that of control eyes by D28. The choroid became thicker during recovery in FD eyes compared to control eyes, in all groups. Choroidal thickness was significantly greater in FD eyes of chickens raised under SL-6500 than in animals raised under FL-4000 (*P* < 0.01). The diameter of scleral collagen fibrils was significantly greater in recovering FD eyes of chickens raised under SL-6500, than in those raised under FL-4000 (*P* = 0.04) and SL-4000 (*P* = 0.002).

**Conclusions:** Compared to fluorescent light, moderate intensities of full-spectrum Sunlike-LEDs can accelerate recovery from form-deprivation myopia in chickens, potentially through choroid-mediated pathways increasing the diameter of scleral collagen fibrils. This study highlights an important implication of the spectral content of white light on ocular growth and emmetropization.

## Background

Myopia is due to a failure in matching the axial length of the eye to the focusing power of its optics, during eye growth, which causes images of distant objects to be focussed in front of the retinal photoreceptors [1]. The prevalence of myopia is steadily rising, and it is estimated that almost 50% of world’s population will be myopic by 2050 [2] making this condition a major socio-economic burden [3]. For instance, in Singapore, 69% of people are myopic by 15 years of age, and approximately US$755 million [4] is spent on optical correction for myopia annually. In East Asia and South East Asia, nearly 95% of the population depends on eyeglasses or contact lenses [5,6]. Although blurred vision can be corrected with glasses, lenses or refractive surgery, there are still risks of blindness from pathologies associated with high myopia [7].

With the strong need to delay myopia onset or slow its progression, several different approaches have been proposed, including increasing time spent outdoors by children [8]. Recent studies have shown that children who spend more time outdoors have a lower risk of developing myopia and experience a reduction in myopia progression [9–13]. Guggenheim *et al*. [14] have shown that the protective effects of outdoor exposure are independent of physical activity, while Donovan *et al*. [15] have found myopia progression to be slower during the summer, possibly because of increased outdoor exposure.

According to Lingham *et al*. [16], the protective effect of outdoor light against myopia is most likely due to one or both of the following factors (which are potentially sub-optimal in indoor lighting): 1) high light intensity and 2) favourable spectral composition of light. Apart from this, circadian rhythm and spatial frequency characteristics of the visual environment are also considered as potential cues; however, there has been limited clinical investigation of this notion [16]. The direct illumination from the sun at noon can rise above 130,000 lux, whilst that in shaded areas outdoors can range from 15,000 to 25,000 lux [17].

Recently, lower levels of light, from 5556 to 7876 lux, have also been found in the shade under trees, while an open-field light intensity during different times of day was reported to range from 11,080 to 18,176 lux in cloudy conditions [18]. In comparison, indoor illumination usually ranges between 100 and 500 lux [17]. The spectral composition of sunlight also differs from that of artificial indoor lighting. Sunlight has a full spectrum that includes wavelengths ranging from ∼300 nm to ∼1200 nm, whilst standard fluorescent indoor lights, for example, emit wavelengths ranging from ∼400 nm to ∼700 nm in a spiked distribution, peaking in blue, red and green [19]. Furthermore, the correlated colour temperature (CCT) of sunlight is dynamic throughout the day, ranging from ∼2,000K at sunrise or sunset to over 10,000K on a clear blue sky at mid-day.

Several experimental studies using animal models have highlighted the importance of the spectral composition of light on axial ocular growth and myopia development [20,21]. For instance, chickens [20], and guinea pigs [22] raised in longer wavelengths of light are more susceptible to ocular axial elongation and increased myopia, while exposure to monochromatic short-wavelength blue lights either induces hyperopia or protects against myopia development, in chicks [21], guinea pigs [23], and some rhesus monkeys [24]. Conversely, exposure to long wavelength monochromatic lights has also been reported to be protective against myopia in rhesus monkeys [25]. To date, the impact of the spectral distribution of artificial white light on refractive error development remains under-investigated.

In this study, we evaluated the impact of moderate intensities of light-emitting diodes (LEDs) with two different CCTs having full, sunlight-like emission spectra, on ocular growth and myopia development and recovery in a chicken model of form-deprivation myopia.

## Methods

### Animals and Experimental Paradigms

One-day-old (D1) male Lohmann Brown chicks were obtained from a hatchery in Malaysia and were raised in a temperature-controlled enclosure (31 ± 1.5°C) with food and water *ad libitum*. Monocular form-deprivation myopia was induced by placing a 3D-printed translucent diffuser over one randomly selected eye. Diffusers were secured around the chicken’s eye using velcro rings, and were inspected twice daily during the 12 h light period to ensure cleanliness. Contralateral eyes remained uncovered and served as controls.

Chicks were reared under a 12:12 hour light/dark cycle (0700-1900h) from day one (D1) until D29. On D1, chickens were randomly divided into three groups based on the spectrum and CCT of isoluminant ambient lighting conditions: Group FL-4000 (n=18) was reared under fluorescent light having a CCT of 4000K (T5 fluorescent tubes, OSRAM GmbH, Munich, Germany); Group SL-4000 (n=12) was reared under a full-spectrum SL LEDs having a CCT of 4000K (T5 Sunlike LEDs, Seoul Semiconductor Co Ltd, Gyeonggi-do, South Korea); and Group SL-6500 (n=9) was reared under a full-spectrum SL LEDs having CCT of 6500K (T5 Sunlike LEDs, Seoul Semiconductor Co Ltd, Gyeonggi-do, South Korea). Light fixtures were controlled by a Helvar DIGIDIM 910 router (Helvar, Dartford Kent, United Kingdom). The spectral composition of the lights (fluorescent and sunlike) were assessed using a calibrated spectroradiometer ILT950 (International Light Technologies, MA, USA). The spectral composition of daylight/sunlight were assessed using CAS 140B-152 spectrometer (Instrument Systems GmbH, Munich, Germany) to compare with the spectrum of Sunlike LEDs. Average light intensities were calibrated at the chicken eye level in all directions of gaze using a calibrated ILT 5000 radiometer (International Light Technologies, MA, USA). (Figure 1)

**Figure 1.**
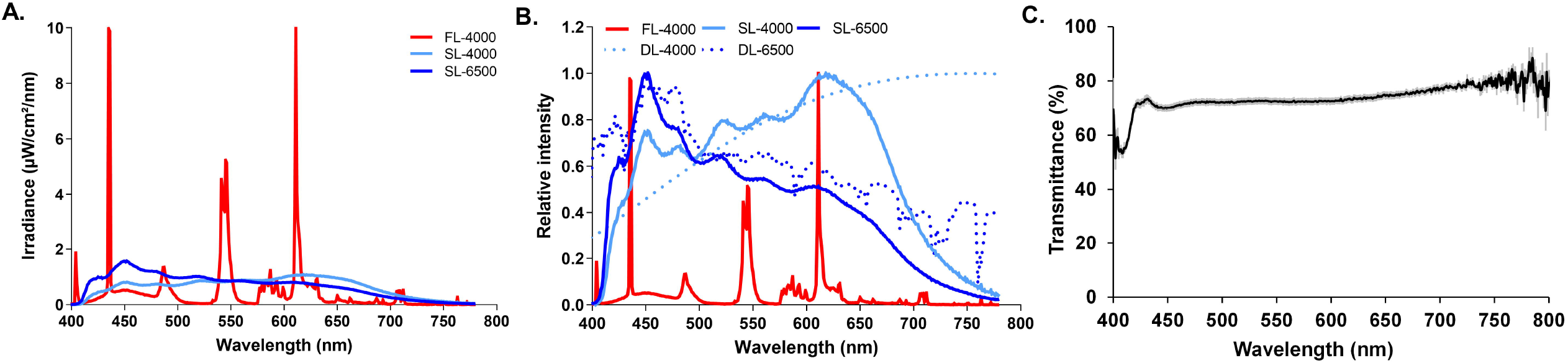
Spectral distribution of fluorescent (4000K: FL-4000), full-spectrum LED (4000K: SL-4000 and 6500K: SL-6500) light-sources, expressed in absolute (A) and relative (B) irradiance. Panel B also includes measurements of daylight/sunlight (4000K: DL-4000 and 6500K: DL-6500) for relative comparison of spectral distribution between artificial and natural sunlight **C**. The average (mean ±SD) light transmittance of 10 diffusers used in this study.

### Ocular measurements *in vivo*

Ocular axial length (AL), non-cycloplegic refraction, choroid and retina thicknesses and anterior chamber depth (ACD) were measured in the three experimental groups on D1, D7, D14, D22 and D28. Baseline ocular parameters were measured on D1 before the application of the diffusers. During the form-deprivation period, the diffusers were removed on D7 and D14 for a brief period to perform ocular measurements. All measurements were carried out on a non-anesthetized

### Axial Length

AL was measured *via* A-scan ultrasonography using a PacScan Plus (SonomedEscalon, NY, USA) at 10 MHz frequency. AL is defined as the distance between the echo spike corresponding to the anterior surface of the cornea and most anterior spike originating from the retina. The median of at least five scans was calculated per eye for each animal.

### Refraction

An automated version of infrared photoretinoscopy was used to measure non-cycloplegic refraction of the chicken’s eye, as previously described by Schaeffel & Howland [26]. Using an adjustable platform, chicks were gently positioned ∼1 meter away from the infrared photoretinoscopy. The median of multiple (3-5) measurements during resting refraction was recorded and utilized in this study.

### Choroidal and retinal thicknesses

Spectral-domain optical coherence tomography (SD-OCT) (Spectralis, Heidelberg Engineering, Inc., Heidelberg, Germany) was used to image the posterior segment of the eye following the protocol adopted by Lan et al. [49]. Choroidal and retinal thicknesses were measured using the Spectralis OCT reviewer software (Heidelberg Eye Explorer, Version 1.10.2.0, Heidelberg, Germany). The average of five measurements per image was utilized in this study.

### Anterior chamber depth

ACD was measured using the RTVue anterior segment OCT (Optovue, Inc., Fremont, CA, USA). An additional corneal lens adapter (cornea anterior module-low magnification [CAM-L]) was attached and the ‘desaturate’ function was enabled for better visualization. Scans of 6 x 6 mm were obtained throughout the study. Imaging was carried out along the nasal-temporal axis.

### Ocular measurements *ex vivo*

To evaluate the impact of study lights on scleral collagen remodelling, animals were euthanized on D29 after completion of the *in vivo* measurements, by administering 2.5mg/kg bodyweight of pentobarbitone sodium (Jurox Pte. Ltd, Rutherford, NSW, Australia) through intraperitoneal injection. Death was further confirmed through cervical dislocation. The eyes were enucleated and scleral strips of 5-10 mm length and 3 mm width were dissected from the posterior pole of all experimental animals. These flat-mounted strips were fixed in 3% paraformaldehyde (Sigma-Aldrich, Merck, St Louis, USA) with 1.5% glutaraldehyde (Ted Pella, inc., Redding, CA, USA) and stored overnight in fixative at 4°C.

### Field Emission Scanning Electron Microscopy (FE-SEM)

For FE-SEM analysis, the fixed scleral strips were dehydrated in 30%, 50%, 70%, 80%, and 90% ethanol for 10 minutes each, followed by 15 minutes in 95% and 100% ethanol and 5 minutes in hexamethyldisilazane (HMDS, Sigma-Aldrich, Merck, St Louis, USA). Samples were left to air-dry overnight in a biosafety cabinet prior to mounting on electron microscope stubs covered with carbon tape. FE-SEM analysis using the Quanta 200F (JEOL – JSM6701F, Lireweg, The Netherlands) was then carried out after sputter-coating with platinum (JEOL JSC-1200 fine coater, Japan) at an accelerating voltage of 15 kV. The fibrous layer of the sclera was imaged. However, this procedure was carried out through the interior surfaces of the scleral strips since residual adhering extraocular muscles interfered with the imaging procedure. Using the micrographs produced by the FE-SEM, average fiber diameter was calculated using ImageJ (National Institutes of Health, USA). Reported values are the average of measurements from 9 randomly selected fibres in the same scleral tissue.

### Data Analysis

Results are presented as average ± standard deviation (SD). Changes in the measured parameters (AL, refraction, choroidal and retinal thicknesses, ACD) of the FD eye over the duration of experimental procedure (D1 to D28) were compared to control eyes (within each experimental group) then expressed as the differences between the FD and the control eye then compared between experimental groups. After confirming normal distribution of the variables, ocular parameters and differences in inter-ocular parameters were compared between eyes within the same group and between groups using a 2-way repeated-measures analysis of variance (2-way repeated measures (RM)-ANOVA) with time and eye or time and group as within- and between-subject factors, respectively. Whole-body weights of animals in different groups were also compared using a 2-way RM-ANOVA. For those comparisons in which the omnibus test reached statistical significance, pairwise multiple comparison procedures were performed using the Holm-Sidak method. Multiple *t*-test was used to analyse the scleral thickness data obtained through FE-SEM. For all statistical tests the level of significance was set at p<0.05, and for post-hoc analysis the Holm-Sidak correction was applied. Statistics were performed using Sigmaplot 14.0 (Systat Software, Inc., San Jose, CA, USA), and plots were drawn using GraphPad Prism 6 (GraphPad Software, La Jolla CA, USA).

## Results

### Characteristics of the experimental lights

Average illuminances, measured in all directions of gaze within the enclosure, were maintained at 281.8 lux (range: 155-727 lux, FL-4000), 284.5 lux (range: 150-711 lux, SL-4000) and 287.9 lux (range: 161-727 lux, SL-6500) throughout the experimental period. The illuminance was highest when the animal was staring directly at the overhead light source. The spectra of SL LEDs (4000K and 6500K) were similar to that of outdoor sunlight/daylight [DL (4000K and 6500K)], being broader and more homogeneously distributed than that of FL-4000, which was discontinuous – having sharp, spike-like peaks at ∼435, 550 and 620 nm (Figure 1A and B). The total energy delivered at these peak wavelengths was much greater for FL-4000 than for the LED sources; however, FL-4000 delivered much higher energy than the LEDs near the λ_max_ of chicken UVS- and SWS-cones (ca. 420 and 455 nm, respectively), but energy comparable to that of the LEDs at the λ_max_ of LWS-cones (ca. 570 nm), and considerably less than that of the LEDs at the λ_max_ of MWS-cones (ca. 510 nm) (Figure 1A) [27]. The average relative spectral transmittance of form-deprivation diffusers at the level of chicken eye was almost the same across the entire visible spectrum, except at the shortest wavelengths (≤425 nm) (Figure 1C).

### Change in body weight of experimental groups

The average weights of the animals raised under both FL and full-spectrum SL lights were similar (*P* > 0.05) throughout the experimental protocol (Supplementary figure S1).

### Impact of full spectrum LEDs on ocular axial length

The axial lengths of form-deprived (FD) eyes of animals raised under all three lighting conditions were significantly greater overall than those of control eyes (FL-4000: [*F* (1,34) = 87.61, *P* < 0.001]; SL-4000: [*F* (1,22) = 19.07, *P* < 0.001] and SL-6500; [*F* (1,16) = 25.75, *P* < 0.001]), and significant differences were seen FD in all groups from D7 to D14 (Figure 2A, B, C). Following the end of form-deprivation (D14), between D14 and D28, FD eyes exposed to SL-4000 and SL-6500 recovered more rapidly from excessive axial elongation than did FD eyes exposed to FL-4000 (Figure 2A, B, C). By D28, the differences between AL of FD eyes in animals raised under SL-4000 (*P* = 0.14) and SL-6500 (*P* = 0.34), compared to control eyes, were no longer statistically significant; whereas in chicks reared under FL, the AL of recovered FD eyes was still 0.91 ± 0.70 mm longer than that of control eyes, on D28 (Figure 2D) (*P* < 0.001).

**Figure 2.**
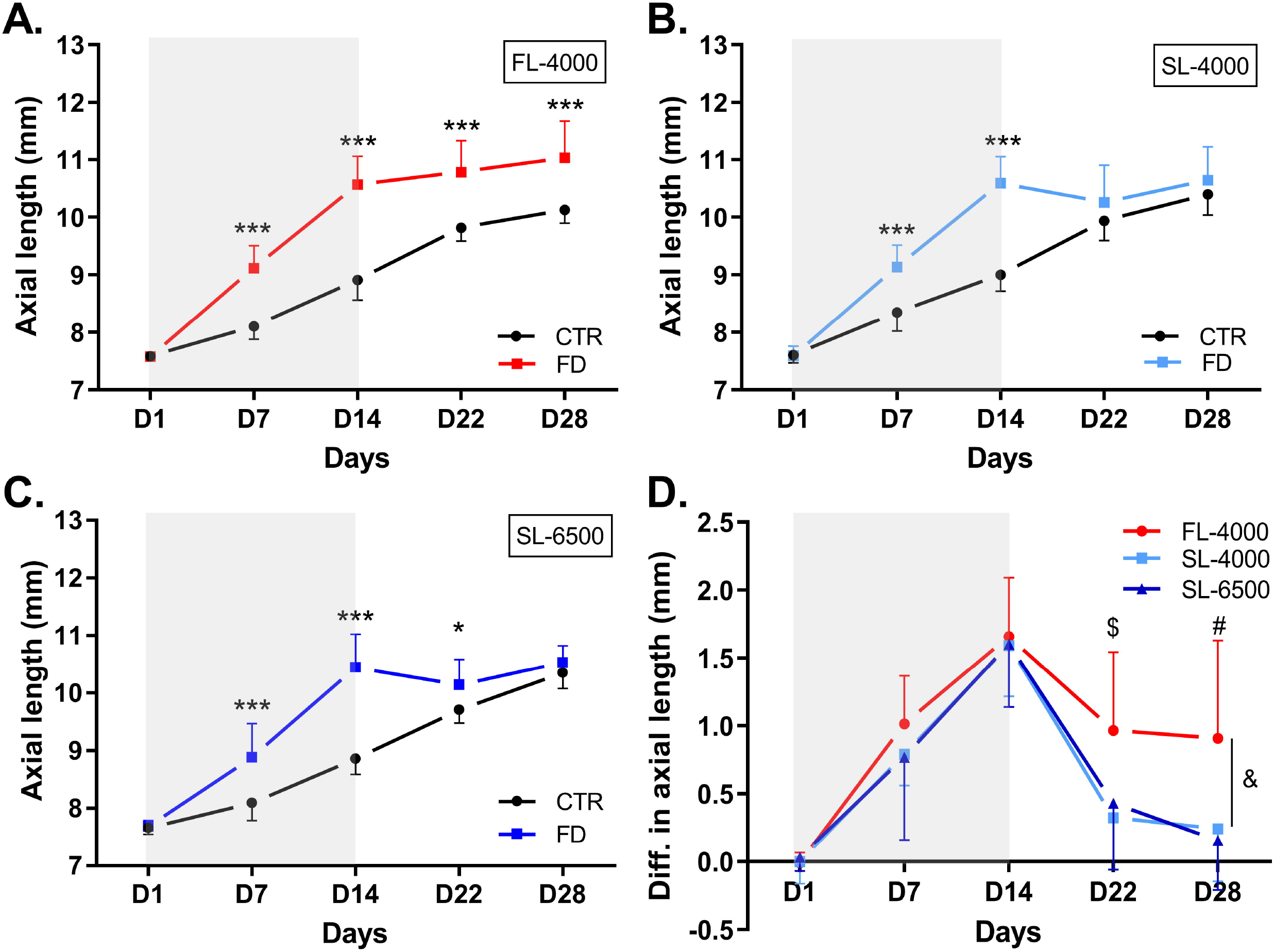
Axial length of FD and control eyes in animals raised under FL-4000 (n = 18) (A), SL-4000 (n = 12) (B), and SL-6500 (n = 9) (C); and inter-ocular differences in axial length of eyes in chicks exposed to FL-4000, SL-4000 and SL-6500 (D). Shaded area between D1 and D14 indicates the period of form-deprivation. &: Significant inter-group difference (*P* = 0.02), for FL-4000 vs both SL-4000 and SL-6500 (both: *P* = 0.04); $: at D22, inter-ocular difference in axial length was significantly higher in FD eyes exposed to FL-4000 than in those exposed to SL-4000 (*P* < 0.001) or SL-6500 (*P* < 0.01); and #: at D28, inter-ocular difference in axial length was significantly higher in FD eyes exposed to FL-4000 than in eyes exposed to SL-4000 (*P* < 0.001) or SL-6500 (*P* < 0.001). *(*P* < 0.05), ***(*P* < 0.001). CTR: Control; FD: Form-deprivation. Data are represented as average ± SD.

The inter-ocular difference in AL was significantly greater overall in the FL-4000 group than in the SL groups ((*F* (2,36) = 4.54, *P* = 0.02; FL-4000 vs SL-4000 (*P* = 0.04), and FL-4000 vs SL-6500 (*P* = 0.049)), but was not significantly different between the SL groups (Figure 2D). This difference was dependent on the time of the experiment (*F* (8,144) = 5.13, *P* < 0.001; viz., on D22 and D28 the average inter-ocular difference in AL of FD eyes exposed to SL-4000 (D22: *P* < 0.001; D28: *P* < 0.001) and SL-6500 (D22: *P* = 0.006; D28: *P* < 0.001) was significantly smaller than in eyes exposed to FL-4000 (Figure 2D). The ALs of control eyes exposed to FL-4000, SL-4000, or SL-6500 were not significantly (*P* > 0.05) different throughout the experimental period.

### Impact of full spectrum LEDs on refractive error

In all groups, the spherical equivalent refractive error of FD eyes exhibited a significant myopic shift compared to control eyes (FL-4000: [*F* (1,34) = 244.44, *P* < 0.001]; SL-4000: [*F* (1,22) = 148.35, *P* < 0.001] and SL-6500; [*F* (1,16) = 29.44, *P* < 0.001]), especially during the form-deprivation period (D7 and D14; Figure 3A-C). Following the end of myopia recovery (D28), the refractions of FD eyes of animals raised under SL-6500 recovered to values similar to those of control eyes (Figure 3C) *(P =* 0.14), whereas the refractions of FD eyes of animals raised under FL-4000 (Figure 3A) (*P* < 0.001) or SL-4000 (Figure 3B) *(P =* 0.04) remained significantly different from those of the fellow control eyes. The inter-ocular differences in refraction between the three groups were not significantly different ((*F* (2,36) = 1.76, *P* = 0.19; FL-4000 vs SL-4000 (*P* = 0.39) and FL-4000 vs SL-6500 (*P* = 0.24)) (Figure 3D). The spherical equivalent refractions of fellow control eyes in chicks exposed to FL-4000, SL-4000 or SL-6500 were not significantly different at any time.

**Figure 3.**
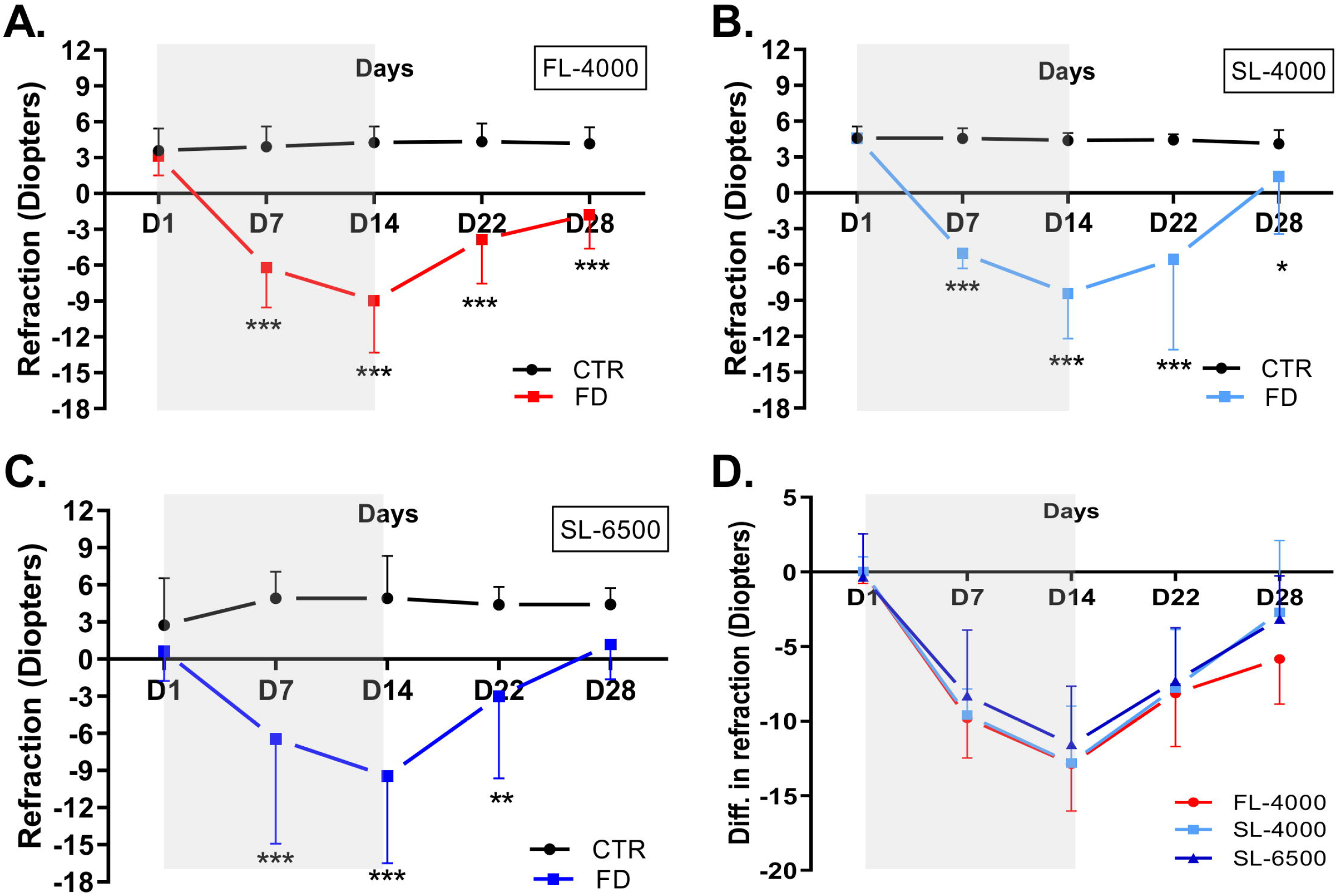
Refractive status of FD and fellow control eyes in animals raised under FL-4000 (n = 18) (A), SL-4000 (n = 12) (B), and SL-6500 (n = 9) (C), and inter-ocular differences in refractive status of eyes exposed to FL-4000, SL-4000 and SL-6500 (D). Statistical significance *(*P* < 0.05) **(*P* < 0.005), ***(*P* < 0.001). CTR-Control and FD-Form-deprivation. Shaded area between D1-D14 time points indicates the period of form-deprivation. Data are represented as average ± SD.

### Impact of full spectrum LEDs on choroidal and retinal thickness measurements

In all groups, the choroidal thickness of FD eyes overall was different from that of fellow control eyes (FL-4000: [*F* (1,34) = 27.63, *P* < 0.001]; SL-4000: [*F* (1,22) = 76.52, *P* < 0.001] and SL-6500; [*F* (1,16) = 145.97, *P* < 0.001]) (Figures 4A, B, C). At specific times, choroidal thickness was greater in FD than in control eyes, on D7 ([FL-4000, *P* = 0.04]; [SL-4000, *P* = 0.02]) and D14 ([FL-4000, *P* = 0.02]; [SL-4000, *P* = 0.03]) (Figure 4A, B); in FD eyes exposed to SL-6500, however, no significant choroidal thinning was observed during the FD period (Figure 4C). After termination of FD at D14, choroidal thickness increased dramatically in the previously-FD eyes exposed to FL-4000, SL-4000 and SL-6500, compared to control eyes (Figure 4A-C), resulting in significantly higher overall inter-ocular differences in choroidal thickness of FD eyes than in the SL-6500 group or the FL-4000 group (*F* (2,36) = 3.92, *P* = 0.03), but no significant differences between choroidal thicknesses in the two SL groups (Figure 4D). Choroidal thickening of the FD eyes under SL-6500 was independent of the time of the experiment. Choroidal thicknesses of control eyes exposed to FL-4000, SL-4000, or SL-6500 were not significantly different at any time.

**Figure 4.**
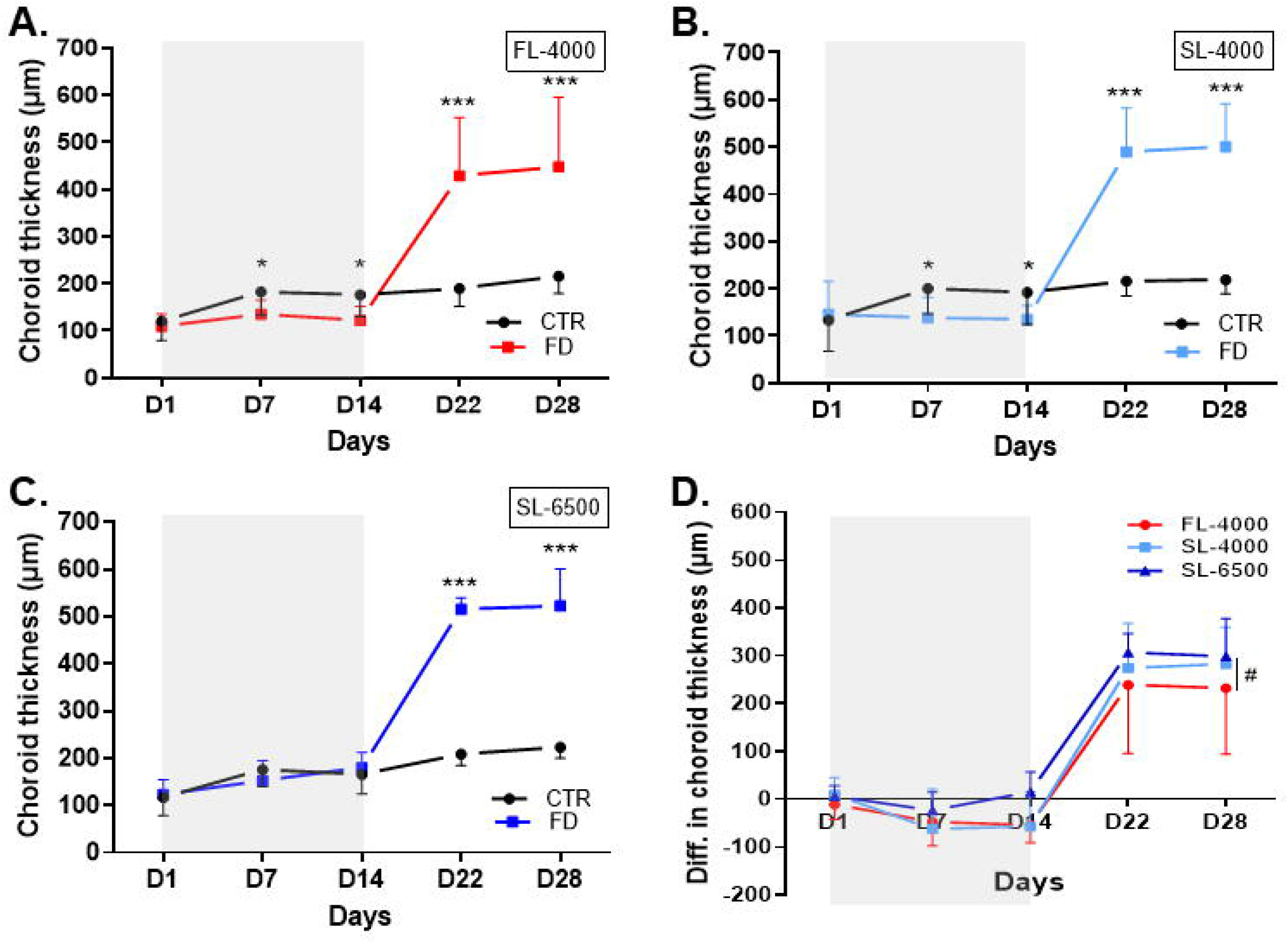
Changes in choroidal thickness of FD and fellow control eyes in animals raised under FL-4000 (n = 18) (A), SL-4000 (n = 12) (B), SL-6500 (n = 9) (C), and inter-ocular differences in choroidal thickness of eyes in chicks exposed to FL-4000, SL-4000 and SL-6500 (D). Statistical significance *** (*P* < 0.001); #: The inter-ocular difference in choroidal thickness in recovering FD eyes exposed to SL-6500 was significantly higher than in those exposed to FL-4000 (*P* = 0.03). CTR: Control; FD: Form-deprivation. Shaded area between D1-D14 indicates the period of form-deprivation. Data are represented as average ± SD.

The retina, too, became significantly thinner by D14 in FD eyes than fellow control eyes, in animals raised under FL-4000, SL-4000 and SL-6500 (FL-4000 [*P* < 0.001]; SL-4000 [*P* < 0.001] and SL-6500 [*P* < 0.001]), while at D7, retinal thinning was found only in the FD eyes of animals raised under FL-4000 (*P* < 0.001) (Supplementary Figure S2A, B). This effect was completely reversed by 8-14 days (D22 and D28, respectively) after diffuser removal on D14, wherein the retinal thickness in FD eyes of all three groups was similar to that in control eyes (Supplementary Figure S2). Inter-ocular differences in retinal thickness in the 3 groups were not significantly different (Supplementary Figure S2D). The retinal thicknesses of control eyes in the 3 groups were not significantly different.

### Impact of full spectrum LEDs on anterior chamber depth

Under all three lighting conditions, ACD was significantly greater in FD eyes than in control eyes (FL-4000: [*F* (1,34) = 24.56, *P* < 0.001]; SL-4000: [*F* (1,22) = 26.96, *P* < 0.001] and SL-6500; [*F* (1,16) = 18.49, *P* < 0.001]) (Figure 5A, B, C). Inter-ocular differences in ACD in the three lighting groups FD were not significantly (*P* > 0.05) different throughout the experimental period (Figure 5D).

**Figure 5.**
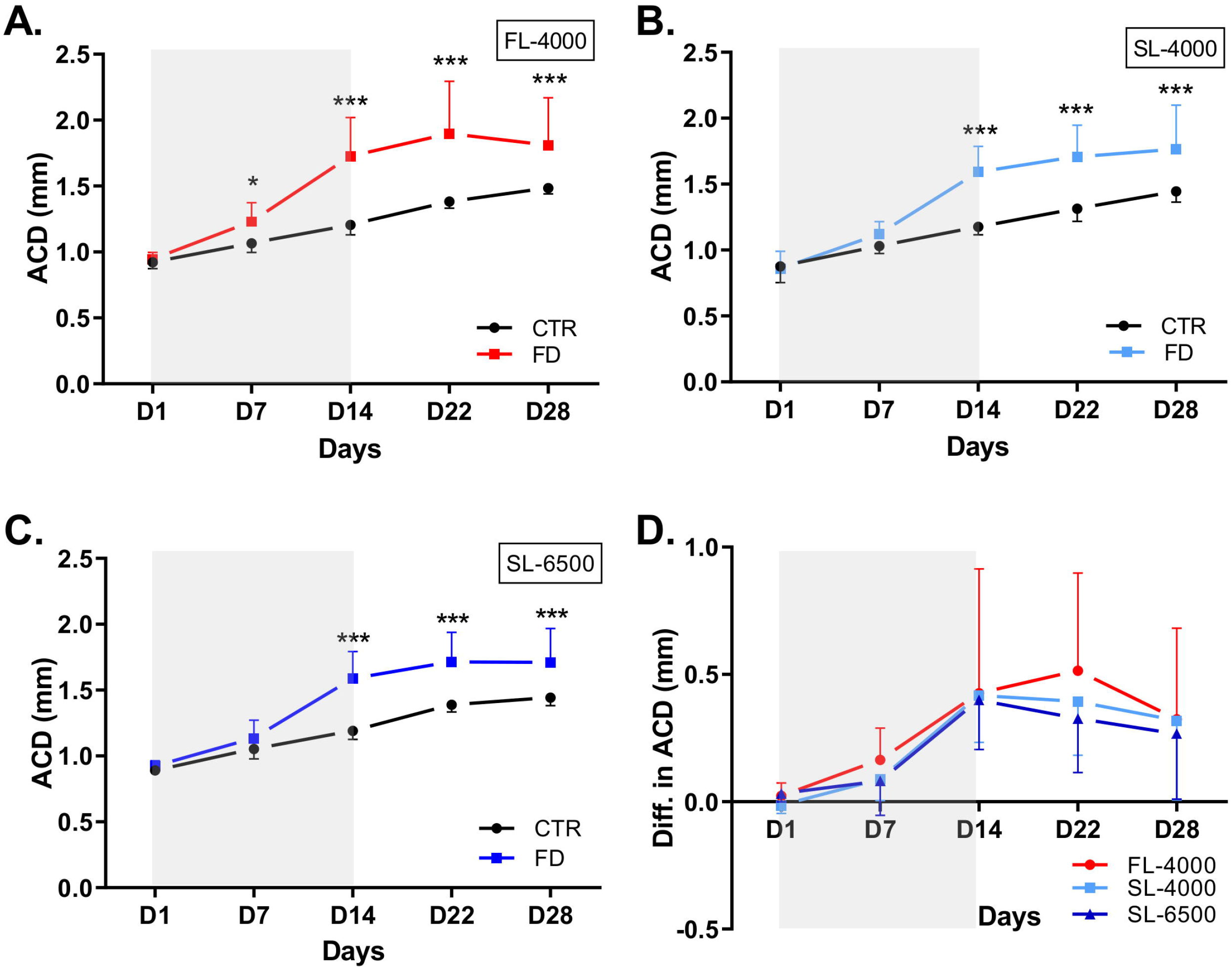
Changes in the anterior chamber depth of FD and fellow control eyes, in animals raised under FL-4000 (n = 18) (A), SL-4000 (n = 12) (B), SL-6500 (n = 9) (C); and inter-ocular differences in ACD of eyes exposed to FL-4000, SL-4000 and SL-6500 (D). ACD: Anterior chamber depth; CTR: Control eyes; FD: Form-deprivation. Shaded area between D1-D14 indicates the period of form-deprivation. Data are represented as average ± SD; ***(*P* < 0.001).

### Scleral collagen fibre remodelling in eyes exposed to SL6500

The FE-SEM analysis reveals that the collagen fibres of recovering FD eyes exposed to SL-6500 were significantly thicker than fibres of recovering FD eyes exposed to FL-4000 (*P* = 0.04) or SL-4000 (*P* = 0.002). The thicknesses of collagen fibre in the control eyes of different study groups were not significantly different (Figure 6A-D). No significant difference (*P* > 0.05) was observed between recovering FD eyes exposed to SL-4000 and FL-4000 (Figure 6E-H).

**Figure 6.**
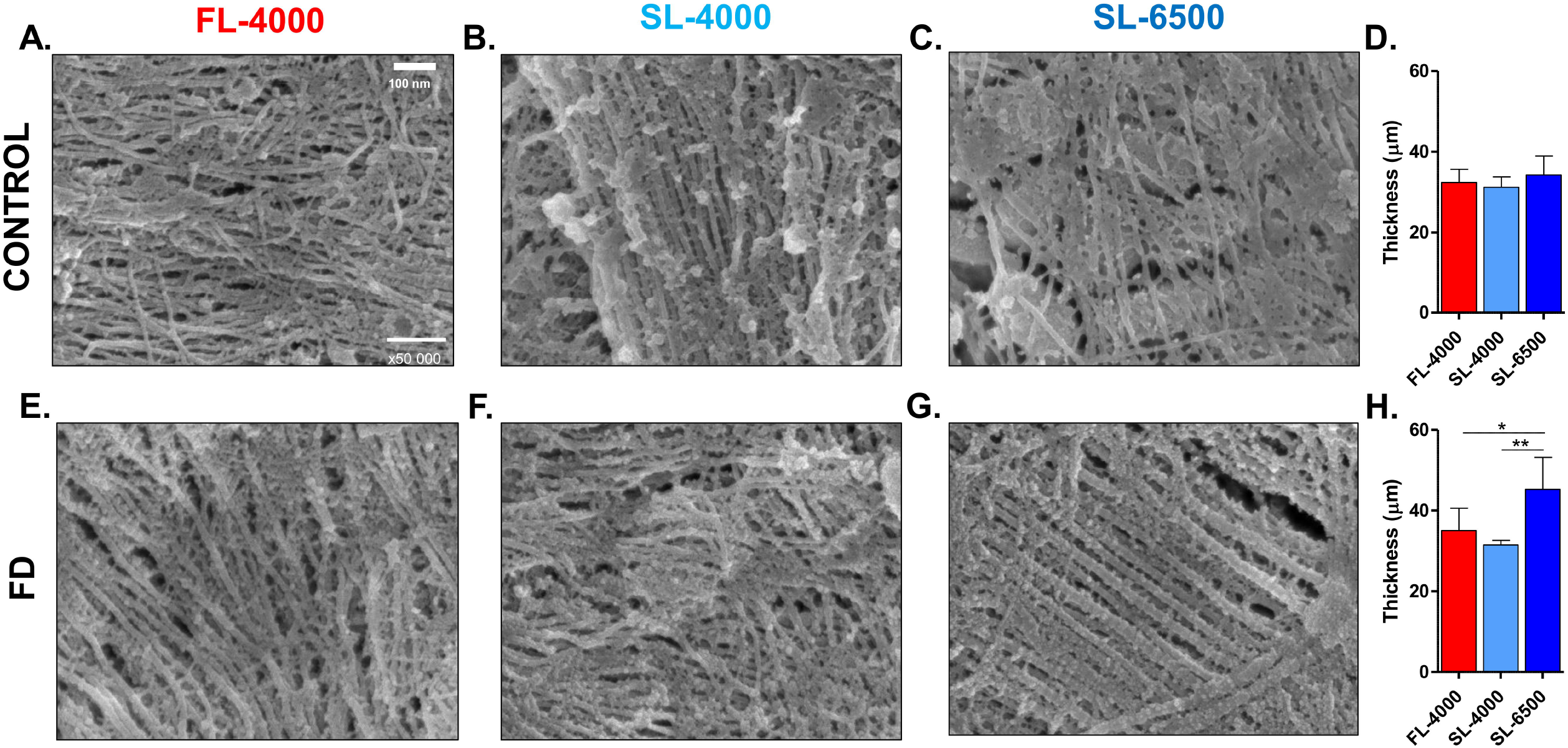
Representative scanning electron micrographs of scleral collagen fibres in the experimental groups (FL-4000, SL-4000 and SL-6500) (A-C and E-G). Average thickness of the scleral collagen fibres from control eyes (D) and FD eyes (H) of different experimental groups (n = 6). *(*P* < 0.05), **(*P* < 0.01).

## Discussion

In this study, we demonstrated that the spectral distribution and CCT of ambient “white” artificial light can affect the recovery from form-deprivation myopia in a chicken model. Irrespective of the CCT, both of the full-spectrum LED lights tested in this study (4000K and 6500K) accelerated recovery from the excessive axial elongation due to form-deprivation, more completely than did isoluminant fluorescent light. None of the lights tested, however, halted the development of form-deprivation myopia, and only the higher-CCT continuous-spectrum LED light (6500K) promoted recovery from myopic refractive error to the extent that there was no significant difference in refraction between the FD and control eyes at the end of the experimental protocol. Furthermore, FD eyes of chicks reared under the higher-CCT continuous-spectrum LED light (6500K) had thicker choroids than FD eyes exposed to fluorescent light, and did not exhibit any choroidal thinning during the development of form-deprivation myopia. In parallel with our anatomical and refraction assessments, *ex vivo* SEM analysis revealed a significant increase in the thickness of scleral collagen fibres in recovering FD eyes exposed to the 6500K light compared to eyes exposed to the other lighting conditions. Whether a causal relationship exists between changes in scleral and choroidal morphology under 6500K light and recovery from FD myopia under this light requires further exploration.

Epidemiological investigations have highlighted the protective effect of spending time outdoors (exposure to sunlight) against human myopia [14]. These findings were complemented by experimental research in animal models. For instance, Ashby et al. [28] demonstrated that chicks fitted with diffusers that were removed daily for 15 minutes to expose under sunlight (∼30,000 lux) significantly reduced the excessive increase in axial length, when compared to 15 minutes exposure to normal laboratory light (∼500 lux). Similarly, exposure of young rhesus monkeys to high-intensity sunlight (average ∼40,000 lux) inhibited the myopic shift induced by monocular -3.0 D lenses as well as the small shift away from hyperopia in fellow control eyes [29]. This protective effect against physiological and experimental myopia might be attributed to any one of many characteristics of sunlight, such as high intensity and full spectral composition [30]. While many investigations have focused on the impact of intense light on experimental myopia development [31–35], only a few have investigated the impact of the spectral characteristics of ambient, moderate intensity, visible white light on refractive error development [19,36]. In our current investigation, the improved recovery from form-deprivation myopia under moderate light levels could be attributed to the spectral content and distribution of Sunlike LEDs, which are similar to those of sunlight. Sunlike LEDs, however, did not stop the development of form-deprivation myopia. These findings are in agreement with Li *et al*. [19], who reported no significant effect of full-spectrum halogen light on refractive error development, in eyes having unrestricted vision compared to eyes having lens-induced myopia. Nevertheless, there are critical differences between our study and that of Li *et al*. [19]. For instance, our study used a form-deprivation myopia model, while the latter used lens-induced myopia. The mechanisms underlying these two myopia models are different in some aspects. [37]. In addition, unlike Li *et al*. [19] we also investigated the effect of moderate levels of full-, continuous-spectrum light during the recovery phase, and the Sunlike LED spectra used in this study were different from that of Halogen light used by Li *et al*. [19].

While both SL-4000 and SL-6500 lights were capable of accelerating recovery from excessive axial elongation, only SL-6500 produced complete recovery from the induced myopic refractive error by 14 days after the termination of form deprivation (day 28 of the experiment). This additional and peculiar impact of SL-6500 could be attributed to its blue-enriched spectrum (Figure 1B), similar to the spectrum of sunlight around noontime. Ocular growth and emmetropization are dependent upon chromatic cues [38], and exposure to monochromatic blue light has been shown to induce hyperopia in some animal models (e.g., chickens [20,21,39] and guinea pigs [22,40]) but not others (e.g., tree shrews [41] and monkeys [25,42]). Previous reports have also suggested a protective effect of blue-enriched white light (CCT > 9000K) against excessive axial elongation in form-deprivation myopia in chickens [36]. Whether this protective effect of blue-enriched light is due to longitudinal chromatic aberration [43] or other phenomena, remains to be clarified. According to Rucker *et al*. [44] the spectral composition of broad-spectrum light, might influence emmetropization and myopia development by retinal mechanisms (circuitry) sensitive either to wavelength per se, or to wavelength-selective defocus due to longitudinal chromatic aberration. However, the emmetropization process can be facilitated by exposure to different visual environment, presenting visual stimuli over broad spectral and temporal ranges and rich in S-cone contrast. These are supposed to be the characteristics of an outdoor lighting environment, mimicked here by Sunlike LEDs.

Choroidal thinning is a particular anatomical change related to myopia, whether in humans [45,46] or animal models [36,47]. In our study, form-deprivation induced choroidal thinning in FD eyes exposed to fluorescent light and SL-4000. However, the choroid was thicker overall in FD eyes exposed to SL-6500 than in those exposed to fluorescent light, and there was no choroidal thinning under SL-6500 during form-deprivation [48]. While it has already been reported that intense light (15,000 lux) induces choroidal thickening in chickens [49], here we have shown that choroidal thickening in healthy and recovering FD eyes is also dependent upon the spectral composition of light. After terminating form-deprivation, the choroids of recovering FD eyes became considerably thicker, irrespective of the lighting condition. This increase in choroidal thickness is considered a compensatory mechanism for the resultant refractive error [47] and has been attributed to changes such as the expansion of choroidal lacunae, increase in choroidal capillary permeability, proteoglycans production, aqueous humor outflow through uveo-scleral routes into the choroid, and decreased tone of the choroidal smooth muscle [50,51].

One of the potential pathways for light-driven myopia-control is through retinal signalling molecules, acting via the retinal pigment epithelium and choroid to regulate scleral structure and biomechanics [52]. In mammals, the sclera encompasses ∼80% of the eye-wall and is recognized as the main load-bearing tissue of the eye; this is achieved through closely packed collagen fibres [53]. Earlier studies on both humans and animals have shown that alterations to collagen micro-architecture and subsequent impact on biomechanical properties of sclera may also contribute to the myopia development [54,55]. In addition, a recent meta-analysis of genome wide association study comprising 160,420 participants of mixed ethnicity (European and Asian) revealed 140 genetic associations linked with light-dependent pathways, which include genes involved in glutamate receptor signalling (*GNB3*) and dopaminergic actions (*DRD1*). These are involved in the light-dependent retina-to-sclera signalling cascade, and therefore are potentially involved in the visual regulation of ocular growth [56]. Furthermore, Kusakari *et al*. [57] demonstrated that diameter of the collagen fibres at the posterior pole of form-deprived eyes were smaller when compared to controls. In the present study, we report that the exposure of recovering FD eyes to high-CCT, full-spectrum SL-6500 light is possibly associated with thicker scleral collagen fibres. Additional studies dedicated to the impact of spectrally tuned light on scleral structure and biomechanics are warranted to better understand light-driven myopia control.

Our study has a few limitations. First, the effect of CCT could not be explained clearly, because we selected LEDs having closely related CCT (4000K and 6500K). These two CCTs, however, were selected to mimic a real-world lighting setup; the impact of higher CCT full-spectrum light mimicking sunlight under a blue sky (∼10,000K) deserves further investigation. Second, we compared the impact of broad-, full-spectrum Sunlike LEDs to that of discontinuous-spectrum fluorescent light. Future studies should compare the effects of Sunlike LEDs to those of standard LEDs used in households. Finally, owing to differences in the ocular systems of chicks and humans, our findings are not directly translatable to understanding and treating human myopia. Future studies, benefitting from insights provided by non-primate animal studies such as ours, should investigate the impact of full-spectrum high CCT lighting on ocular growth in primate models.

## Conclusion

This study, using the chicken model of form-deprivation myopia, reveals that moderate levels of continuous-, full-spectrum LED lights that mimic sunlight can promote recovery from myopia after restoring normal viewing; and that full-spectrum light with higher CCT can prevent the choroidal thinning that accompanies myopia. Our findings provide evidence that the spectral composition of indoor light could affect ocular growth and and emmetropization, and they open new research avenues for light-centred, passive myopia-control.

## List of abbreviations

ACDw: Anterior chamber depth
AL: Axial length
CAM-L: Cornea anterior module-low magnification
CCT: Correlated colour temperature
CTR: Control
DL: Daylight/sunlight FD
FD: Form-deprived
FE-SEM: Field Emission Scanning Electron Microscopy
FL: Fluorescent
HMDS: Hexamethyldisilazane
LEDs: Light-emitting diodes
RM-ANOVA: Repeated measure-analysis of variance
SD: Standard deviation
SD-OCT: Spectral-domain optical coherence tomography
SL: Sunlike

## Declarations

### Ethics approval and consent to participate

All animals used in this study were treated in accordance with the ARVO Statement for the Use of Animals in Ophthalmic and Vision Research. Protocols used were approved by the SingHealth Institutional Animal Care and Use Committee (IACUC; AAALAC Accredited; 2018/SHS/1444).

### Consent for publication

Not applicable

### Availability of data and materials

The datasets used in the current study are available from the corresponding author on request.

### Authors contributions

R.P.N. designed research; A.R.M., S.L.W.Y. and L.Y.C. performed research; A.R.M., R.P.N. and S.L.W.Y. analysed data; A.R.M. and S.L.W.Y wrote the paper; R.P.N., S.M.S., V.A.B. and D.M. provided critical inputs; all authors reviewed the paper and R.P.N., acquired funding.

## Acknowledgements

We would like to thank Prof. William Stell for his advice and for reviewing this manuscript. We also thank Dr. Royston Tan for 3D printing the chicken diffusers and Mr. Noel Sng for helping us in designing the chicken enclosure.

## Figure Legends

**Supplementary Figure S1.** Average weight of animals in experimental groups raised under FL-4000 (n = 18), SL-4000 (n=12) and SL-6500 (n = 9) throughout the experiment. Shaded area between D1 and D14 indicates the period of form-deprivation. Data are represented as average ± SD. Body weights at all durations were not significantly different, and were not affected by FD or recovery (P>0.05).

**Supplementary Figure S2.** Changes in the retinal thickness in FD and fellow control eyes in animals raised under FL-4000 (n = 18) (A), SL-4000 (n = 12) (B), SL-6500 (n = 9) (C); and inter-ocular differences in retinal thickness in eyes exposed to FL-4000, SL-4000 and SL-6500 (D). CTR: Control; FD: Form-deprivation. Shaded area (D1-D14) indicates the period of form-deprivation. Data are represented as average ± SD; ***(P < 0.001).

