## Supplementary figures and images for "Recovery from form-deprivation myopia in chicks is dependent upon the fullness and correlated colour temperature of the light spectrum"

### S1.tif

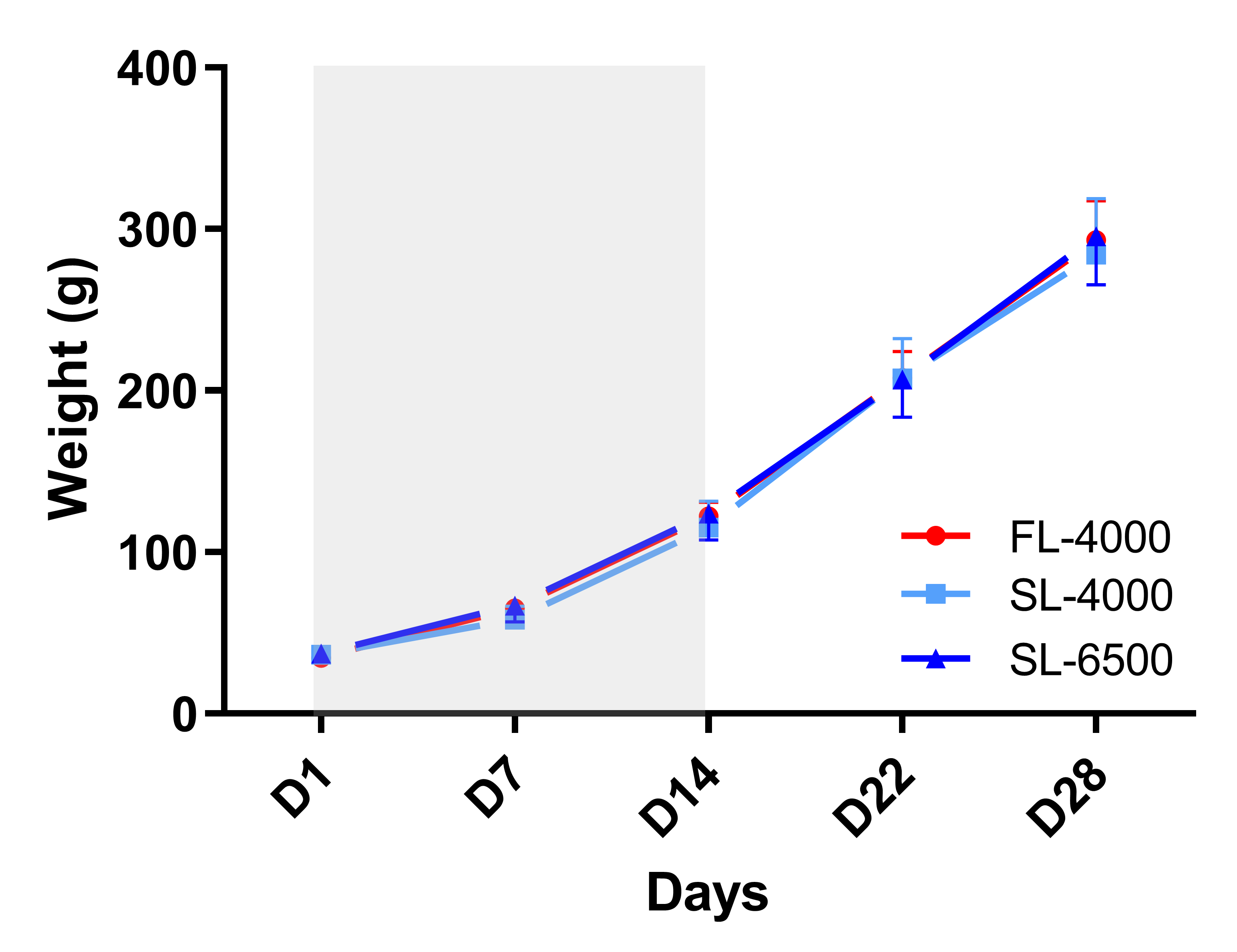

### S2.tiff

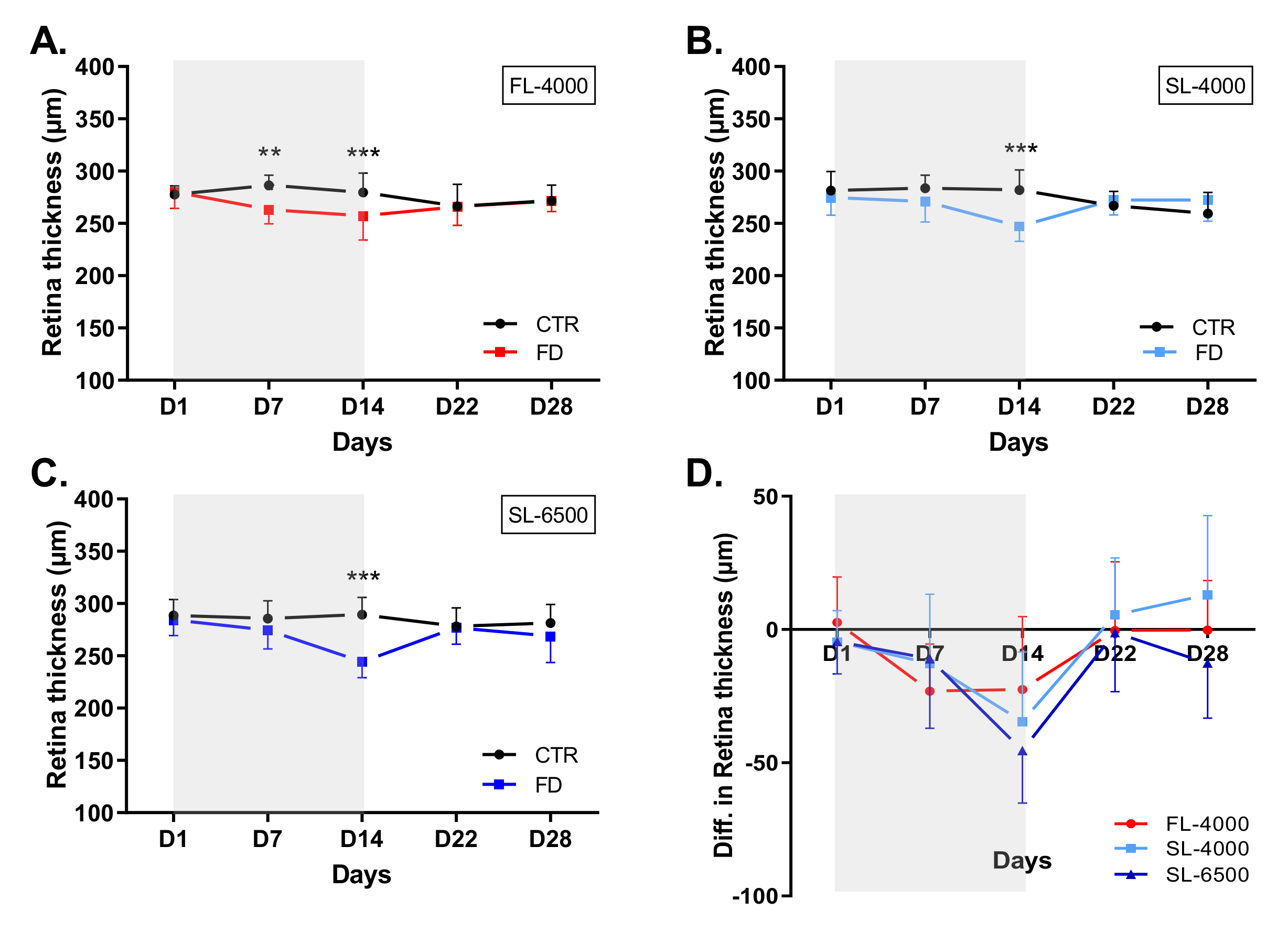
